# The floral development of the allotetraploid *Coffea arabica* L. correlates with a small RNA dynamic reprogramming

**DOI:** 10.1101/2023.08.23.554532

**Authors:** Thales Henrique Cherubino Ribeiro, Patricia Baldrich, Raphael Ricon de Oliveira, Christiane Noronha Fernandes-Brum, Sandra Marisa Mathioni, Thaís Cunha de Sousa Cardoso, Matheus de Souza Gomes, Laurence Rodrigues do Amaral, Kellen Kauanne Pimenta de Oliveira, Gabriel Lasmar dos Reis, Blake C. Meyers, Antonio Chalfun-Junior

**Affiliations:** Laboratory of Plant Molecular Physiology, Plant Physiology Sector, Department of Biology, Federal University of Lavras (UFLA), MG 37200-900, Brazil; Donald Danforth Plant Science Center, Saint Louis, MI 63132, USA; Laboratory of Bioinformatics and Molecular Analysis, Federal University of Uberlandia (UFU), Campus Patos de Minas, MG 38700-128, Brazil; Division of Plant Sciences and Technology, University of Missouri-Columbia, Columbia, MI 65211, USA

**Keywords:** *Coffea arabica*, Floral development, microRNA, phasiRNAs, smallRNAs

## Abstract

Non-coding and coding RNAs are key regulators of plant growth, development, and stress responses. To investigate the types of transcripts accumulated during the vegetative to reproductive transition and floral development in the *Coffea arabica* L., we sequenced small RNA libraries from eight developmental stages, up to anthesis.
We combined this data with messenger RNA and PARE sequencing of two important development stages that marks the transition of an apparent latent to a rapid growth stage. In addition, we took advantage of multiple *in silico* tools to characterize genomic loci producing small RNAs such as phasiRNAs, miRNAs and tRFs.
Our differential and co-expression analysis showed that some types of small RNAs such as tRNAs, snoRNAs, snRNAs and phasiRNAs preferentially accumulate in a stage-specific manner.
Members of the miR482/miR2118 superfamily and their 21-nucleotide phasiRNAs originating from resistance genes show a robust co-expression pattern that is maintained across all the evaluated developmental stages. Finally, the majority of miRNAs accumulate in a family-stage specific manner, related to modulated hormonal responses and transcription factors expression.

**Societal Impact Statement:** This research holds potential to benefit millions of coffee-producing families in over 60 countries. We uncovered molecular regulatory mechanisms governing flower development, one of the causes for the *Coffea arabica*’s uneven ripening. The absence of uniformity in coffee production, spanning from floral induction to branch senescence, has a detrimental impact on the final product’s quality. These insights will inform strategies for controlled coffee maturation, leading to improved, uniform harvests.

## Introduction

In polyploid plants like *Coffea arabica* L., which provides the basis for much of the world’s coffee consumption, RNAs are transcribed from co-existing genome versions within a cell (Scalabrin *et al*., 2020). Each version of its genome comes from one ancestral progenitor from an intraspecific cross between two ancient specimens, *Coffea eugenioides* Moore and *Coffea canephora* Pierre ex Froehner (Lashermes *et al*., 1999). The *C. arabica* allopolyploidy (2*n* = 4*x* = 44) may be beneficial by improving transcriptional homeostasis relative to its diploid parents (Bertrand *et al*., 2015). Nevertheless, allopolyploidy can add complexity in the meiotic cell division (Lloyd & Bomblies, 2016).

The non-uniformity in *C. arabica* flowering and ripening process is a serious economic problem that affects its production and beverage quality, even more so under climate change (Davis *et al*., 2019; de Oliveira *et al*., 2020). This lack of uniformity is in part due to the biennial phenological cycle of coffee (Camargo & Camargo, 2001) that comprehends periods of vegetative meristem’s formation and floral buds induction/development (de Oliveira *et al*., 2014). In addition, being perennial, it must keep the shoot apical meristem in a vegetative state to allow simultaneous growth and reproductive development (Camargo & Camargo, 2001). Although some endogenous and environmental factors are known to be involved in the vegetative to reproductive transition of coffee (López *et al*., 2021), the role of small RNAs and non-coding RNAs is a topic overlooked. To fill this knowledge gap, we conducted a comprehensive analysis of RNA accumulation throughout the flower differentiation stages in *C. arabica*.

Here we evaluated messenger RNAs (mRNAs), microRNAs (miRNAs), phased small interfering RNAs (phasiRNAs), small nuclear RNAs (snRNAs), small nucleolar RNAs (snoRNAs) and transfer RNAs (tRNAs). We found that different types of small RNAs (sRNAs) are preferentially accumulated in a stage-specific manner. Furthermore, our analysis provided valuable insights into the interactions of these RNAs with molecular machinery, including hormone crosstalk and defense response systems.

Finally, we detected sRNA accumulation changes in two contrasting development stages often anatomically classified as a single stage (Morais *et al*., 2008). Buds ranging from 3 mm to 6 mm, known as G4, are commonly labeled as dormant or latent. Despite their months-long apparent anatomical latency, our transcriptional analysis reveals the accumulation of 21 and 24-nt phasiRNAs. Following environmental cues like prolonged drought followed by rain, phasiRNA levels decrease sharply, while snoRNAs and tRNAs experience a rapid but brief increase. This finding led us to propose novel transcriptomic-based classification that adapts the phenology classification of Morais *et al*. (2008) to discriminate buds > 3 mm and < 6 mm into two new stages: an early stage now called S3 – transcriptionally characterized by the accumulation of 24-nt phasiRNAs and the occurrence of meiosis (Pimenta de Oliveira *et al*., 2023) – and a late stage called S4, characterized by the fast accumulation of tRNAs, snoRNAs and the resumption of developmental programs.

## Materials and Methods

### Plant material

To obtain data for RNAseq, small RNAseq (sRNAseq) and Parallel Analysis of RNA Ends (PARE), we selected 5-year-old *C. arabica* plants of two cultivars; “Siriema VC4” and “Catuaí Vermelho IAC 144”. These plants were grown at the experimental field in the Federal University of Lavras (UFLA), Brazil (21°13’ S, 44°58’ W) and were maintained with standard cultivation practices (Vieira, 2008). After harvesting, all samples were immediately frozen in liquid nitrogen and stored at - 80 °C until total RNA extraction. Sample details are provided in Table S1.

### Library preparation and sequencing

We isolated total RNA with PureLink® Plant RNA Reagent (Invitrogen). Then, for the sRNAseq libraries, we performed the size selection using denaturing Urea-PAGE gels and library construction using the TruSeq Small RNA Library Preparation kit following the protocol described by Mathioni *et al* (2017). Sequencing of 256 million single-end reads was performed at the University of Delaware Sequencing and Genotyping Center using an Illumina HiSeq 2500 sequencer. For the RNAseq of two development stages (S4 and S5) the libraries were prepared with Illumina TruSeq stranded RNAseq preparation kit. The sampling for the RNAseq libraries was the following; 2 biological replicates x 2 development stages (S4 and S5) x 2 cultivars, rendering 8 strand-specific single-end libraries with a total of 376 million reads. Finally, PARE libraries for S4, S5, pre-meiotic, meiotic and post-meiotic samples were constructed using the protocol described by German *et al* (2009) totaling 123 million reads. The pre-meiotic, meiotic and post-meiotic stages were previously characterized in coffee by Pimenta de Oliveira (2023).

### Identification of miRNAs

We identified conserved mature and precursor miRNAs in the *C. arabica* genome using the procedure described by de Souza Gomes *et al* (2011). The pipeline encompasses stages such as filtering for GC content between 20% and 65%, minimum free energy below −20 kcal/mol, and selecting mature sequences with over 85% identity to plant mature miRNA registered in miRbase Release 22.1(Kozomara *et al*., 2019). Subsequently, we augmented this collection of miRNAs with novel and conserved miRNAs predicted using miRador (Hammond *et al*., 2021) with the most up-to-date criteria to accurately identify plant miRNAs (Axtell & Meyers, 2018). Finally, we developed custom Python scripts to merge miRNAs with identical mature sequences predicted by both methodologies.

### Identification of *PHAS* loci

We mapped ∼1.2 billion quality-controlled sRNAseq reads from 8 stages ranging from nodes containing undetermined cells to flowers in addition to pre-meiotic, meiotic and post-meiotic anthers in the *C. arabica* Caturra genome (Johns Hopkins University, 2018) with Bowtie (Langmead *et al*., 2009). No gaps or mismatches were allowed. The resulting files were processed with ShortStack (Axtell, 2013) to predict phased locus. Next, we manually assessed predicted loci using criteria such as score, length, protein or TE overlap, strand alignment ratio and complexity. Lower complexity values (near 0) signified loci dominated by a few small RNAs, while higher values (near 1) indicated diverse small RNA sets. The alignments were inspected with the Integrative Genome Viewer (IGV). That way, we categorized the 803 candidate PHAS loci into true PHAS loci, Long 24-PHAS-like loci (L24P-like), or, if the locus characteristics were unclear, an unknown type (UNK).

### Differential accumulation analysis of phasiRNA, tRNAs, snoRNA, and snRNAs

To assess the accumulation profiles of sRNA types, we manually compiled a reference FASTA file. This file included non-redundant mature miRNA sequences, all predicted PHAS/PHAS-like loci, as well as 307 representative sequences of non-redundant tRNAs, 157 snRNAs, and 1,203 snoRNAs. Next, we mapped the ∼1.2 billion quality controlled sRNAseq reads from nodes containing undetermined vegetative cells (S0) to flower using Bowtie (Langmead *et al*., 2009) with parameters -k 1 and -v 0 (no mismatches allowed). We conducted differential analysis using edgeR (Robinson *et al*., 2010) considering a small RNA-producing locus as differentially accumulated if its false discovery rate (FDR) < 0.05 and fold change was ≥ 2.

### sRNA co-expression network

To infer co-expression modules (clusters) of sRNAs, we applied procedures from the Weighted Gene Co-expression Network Analysis (WGCNA) R-package (Langfelder & Horvath, 2008). We used the same count data as in the differential analysis, with necessary transformations. Initially, we normalized counts to Counts Per Million (CPM) and then applied a log2 transformation. Subsequently, we computed a Pearson correlation adjacency matrix for all pairs of transcribed small RNAs, powered by ! (soft threshold) set to 6, following the scale-free topology criterion (Zhang & Horvath, 2005). After that, to minimize the effect of noise and spurious associations, we transformed the adjacency matrix into a Topological Overlap Matrix (TOM). Next, a dendrogram, with the co-expression modules as its branches, was inferred based on the average dissimilarity of the TOM using the dynamic tree cut method. Finally, we analyzed the individual co-expression modules with the R package igraph (Csardi & Nepusz, 2006).

### Target prediction

To validate the siRNA targets profiled by Parallel Analysis of RNA Ends sequencing (PARE) we used sPARTA (Kakrana *et al*., 2014) with parameters --map2DD, --validate, -minTagLen 18 and - tarScore N. These degradome analyses were performed to identify genic and intergenic cleavage sites of mature miRNAs and genic cleavage sites of tRNA Fragments (tRFs). The target cutoff score was set to ≤5 for miRNA targets and, more restrictively, ≤3 for tRFs.

The tRFs used for this PARE analysis were processed as follows; First, all quality controlled sRNAseq reads were mapped to the tRNA reference available at the NCBI (Johns Hopkins University, 2018). All fragments mapped without mismatches were then selected based on their length (minimum of 18 and maximum of 24 nt) and expression in CPM (minimum of 1CPM). Doing so, we selected 7,458 tRFs that were used by sPARTA to identify putative cleavage sites.

### Differential expression analysis of protein coding genes

Approximately 376 million single-end RNAseq reads from the stages S4 and S5 were sequenced in eight libraries: four biological replicates for each stage of “Siriema VC4” and “Catuaí Vermelho IAC 144” cultivars. After quality control with trimmomatic (Bolger *et al*., 2014) v.0.33 (parameters ILLUMINACLIP:./adapters:3:25:6 SLIDINGWINDOW:4:28 MINLEN:30) 272 million reads were mapped to the genome with STAR aligner v.2.7.1 (Dobin *et al*., 2013). Then, we quantified the reads uniquely mapped to exons in the genome using the htseq-count script (Anders *et al*., 2015). We carried out differential expression analysis with edgeR (Robinson *et al*., 2010) and a given protein coding gene was deemed differentially expressed (DE) between stages if its FDR was less than 0.05 and fold change ≥ 2. Then, we searched for enriched gene ontology (GO) terms using the online tool agrigo v.2 (Yan *et al*., 2017).

## Results

### A higher number of miRNA precursor loci in *C. arabica* compared to Arabidopsis thaliana reflects 100 million years of divergence

A total of 557 candidate miRNA precursor loci from 296 miRNA families were found in the *C. arabica* genome (Fig. 1 - First inner cycle). Those precursors produce 447 nonredundant mature miRNAs. We discovered that 45 of those precursors generate mature miRNAs representing putative novel family members (Fig. 1 - First inner cycle, violet points). A total of 205 miRNA precursors were found to be encoded in the *C. eugenioides* sub-genome and 263 from the *C. canephora* sub-genome (Fig. 1 - First inner cycle, black dots). Additionally, 89 miRNA precursors were found in the unplaced contigs.

**Figure 1.**
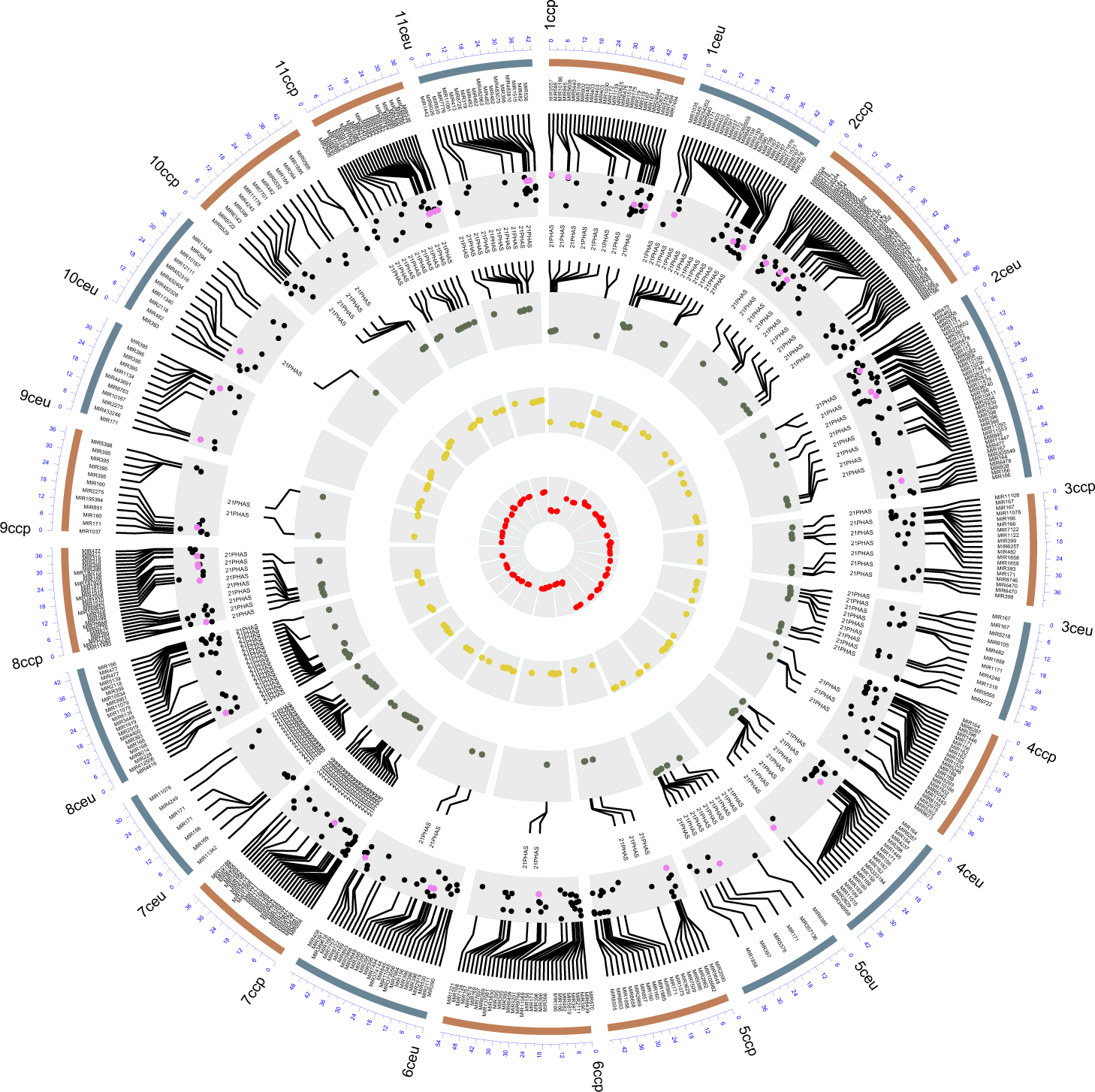
Genome-wide distribution of MIRs, PHAS and PHAS-like loci in the allopolyploid Coffea arabica. The outermost cycle represents chromosomes from each parental ancestor; from C. eugenioides (ceu) or C. canephora (ccp). First inner cycle; Chromosomal coordinates of miRNA precursors, black dots are conserved families while violet dots are putative novel. Second inner cycle; green dots point out the chromosomal coordinates of 21-PHAS loci. Third inner cycle; point out the chromosomal coordinates of 24-PHAS loci. Innermost cycle; red dots point out the chromosomal coordinates of 24-PHAS-like loci.

Among the conserved miRNA families in the Asterids, miR160 was only identified in the *C. canephora* sub-genome whereas the miR828 family is only found in the *C. eugenioides* sub-genome. The miR173 family, known in *A. thaliana* for triggering the biogenesis of both trans-acting small interfering RNAs (tasiRNAs) *TAS1* and *TAS2* (Allen *et al*., 2005), is not present in the *C. arabica*. The families having the greatest numbers of loci were miR482, miR395, miR169, miR167 and miR171 (Table S2) whereas the most accumulated families were, in decreasing order, miR166, miR396, miR482, miR319 and miR8155 (Fig. S1). *A. thaliana* has no miR482 (Zhu *et al*., 2013), which is the most abundant in terms of the number of loci in *C. arabica*. However, Arabidopsis still encodes other abundant loci in *C. arabica* such as miR395, miR169, miR167 and miR171 (Fahlgren *et al*., 2010).

The expansion of the number of miRNA loci in *C. arabica* can be explained by the net rate of flux (birth-death). In the *Arabidopsis* lineage, this rate was estimated to be a net gain of 1.2 to 3.3 *MIR* genes per million years (Fahlgren *et al*., 2010). According to miRbase v. 22 (Kozomara *et al*., 2019) there are 326 miRNA precursors in the *A. thaliana* genome (Kozomara *et al*., 2019). It is estimated that *Coffea canephora* diverged from *Vitis vinifera*, a basal rosid, about 114 to 125 million years ago (Wikström *et al*., 2001; Guyot *et al*., 2012). This suggests that *C. arabica* and Arabidopsis had a shared ancestor around the same period as *C. canephora* and *V. vinifera*. Over the course of 100 million years of evolution and the emergence of allopolyploid *C. arabica*, it’s conceivable that hundreds of microRNA genes were gained and lost, resulting in the observed difference of 231 miRNA precursors between *C. arabica* and *A. thaliana*.

### The majority of 21-*PHAS* loci correspond to disease resistance proteins triggered by the miR482/2118 superfamily

PhasiRNAs in plants arise from long noncoding RNAs (lncRNAs) or protein-coding transcripts. After a miRNA-guided precise cleavage, secondary siRNAs are produced through the action of DICER-like enzymes. We used sRNAseq libraries from vegetative and reproductive organs to identify loci producing phasiRNAs. We identified 173 21-*PHAS* loci, most of them from disease resistance (R) genes containing Nucleotide Binding Leucine-Rich Repeats domains (NB-LRR; Fig. 1 - second internal cycle, green dots; Table S2). PARE analysis shows miRNA triggering at least 51 21*-PHAS*, including 23 triggered by miR482 family members. In addition, 15 21*-PHAS* loci were found to be triggered by a putative novel miRNA, the candidate miR245889. Similarity based analysis suggests that this candidate diverged from the miR482*/*miR2118 superfamily (Fig. S2).

Our prediction of phasing loci allowed us to identify two candidate tasiRNA. Their miRNA target sites were similar to miR390 and miR828. Further analysis of those loci revealed that they are the respective orthologs of *TAS3* and *TAS4* (Fig. S3). We did not find any orthologs for *TAS1* and *TAS2* in accordance with the lack of their conserved trigger, miR173. We also identified two *DICER-like 2* (*DCL2*), one in chromosome 6e and another in 9c, as loci generating phasiRNAs. This phasing of *DCL2* has been described as being triggered by miR1507 in Fabaceae (Zhai *et al*., 2011), and miR6026 or miR10533 in Solanaceae (Baldrich *et al*., 2022). Of these known *DCL2* triggers we could only identify a putative miR6026 of 22 nucleotides being encoded exclusively by the *C. eugenioides* sub-genome. Nevertheless, its accumulation couldn’t be detected in any assessed developmental stage. This suggests the possibility of an as-yet-unknown sRNA with a precision cleavage mechanism enabling the biogenesis of 21-nt phasiRNAs from *DCL2* transcripts.

### 24-PHAS loci are not triggered by any expressed miRNA

Recent reports have highlighted the existence of 24-nt phasiRNAs in numerous eudicots. These small RNAs exhibit significant enrichment in reproductive organs, particularly anthers. (Xia *et al*., 2019; Pokhrel *et al*., 2021; Pokhrel & Meyers, 2022). Because several phased siRNA annotation methods can frequently mistake heterochromatic siRNAs with 24*-PHAS* loci (Polydore *et al*., 2018) we manually evaluated all putative *PHAS* loci with IGV. Doing so, we were able to identify 189 24-*PHAS* loci (Fig. 1 - third internal cycle with yellow dots; Table S3). Of those loci, 56 overlap with annotated protein coding genes (PCG). In addition, 58 of those 24-*PHAS* overlapped with transposable elements (TEs) such as Gypsy (34%) and hAT (22%).

Surprisingly, we did not identify any evident miRNA triggers for 24-nt phasiRNAs. Canonically, miR2275 serves as the common trigger for 24-nt phasiRNA biogenesis (Xia *et al*., 2019). However, it is known to be absent in some lineages such Brassicales, Caryophyllales, Cucurbitaceas, Fabales and Lamiales (Xia *et al*., 2019). In *C. arabica,* we found four putative miR2275 precursor loci, but without evidence of accumulation in the analyzed organs. A similar scenario was previously reported in Solanales like tomato and petunia, where abundant 24-nt reproductive phasiRNAs were discovered in meiotic anthers, yet no identifiable miRNA trigger was found (Xia *et al*., 2019).

### Hundreds of loci demonstrate a 24-nt phasing pattern but excluded as 24-*PHAS* loci

There were 175 long 24-*PHAS*-like loci (L24P-like; Fig. 1 - Fourth internal cycle with red dots; Table S4). We named them L24P-like for multiple reasons, these loci met our phasiRNA locus filtering criteria and primarily originated from non-repetitive genome regions. Like 24-*PHAS* loci, they lacked apparent triggers. However, their length is atypical (Fig. S4), with a median of 10.3 kb for L24P-like compared to around 1.8 kb for 24-*PHAS* loci precursors.

These L24P-like loci resemble “siren” (small-interfering RNA in endosperm) loci in terms of their length and stage specificity (Fig. 2a). Siren loci generate 24-nt siRNAs, first identified for their abundance in rice endosperm (Rodrigues *et al*., 2013; Grover *et al*., 2020). These loci map predominantly to genic and intergenic regions rather than transposable elements (Burgess *et al*., 2022). The siren loci also correspond to approximately 200 loci in *Brassica rapa* and are present in diverse other angiosperms (Rodrigues *et al*., 2013; Grover *et al*., 2020). They are transcribed in maternal organs and thought to induce DNA methylation in filial tissues, thus establishing epigenetic marks in subsequent generations (Grover *et al*., 2020). Siren loci are typically larger than other siRNA-producing loci, and their derived siRNAs are more likely to have unique genome mappings compared to other siRNA categories (Grover *et al*., 2020). In addition, they tend to represent more than 90% of the accumulated siRNAs in developing seeds (Grover *et al*., 2020). In accordance with the patterns of these sirenRNAs, the number of multi-mapped reads of *C. arabica* L24P-like (mean = 14,581, sem = 1,201) are substantially higher than the 24-*PHAS* loci (mean = 8,353, sem = 1,586; two-tailed t-test unequal variance p = 1.1E^-3^).

**Figure 2.**
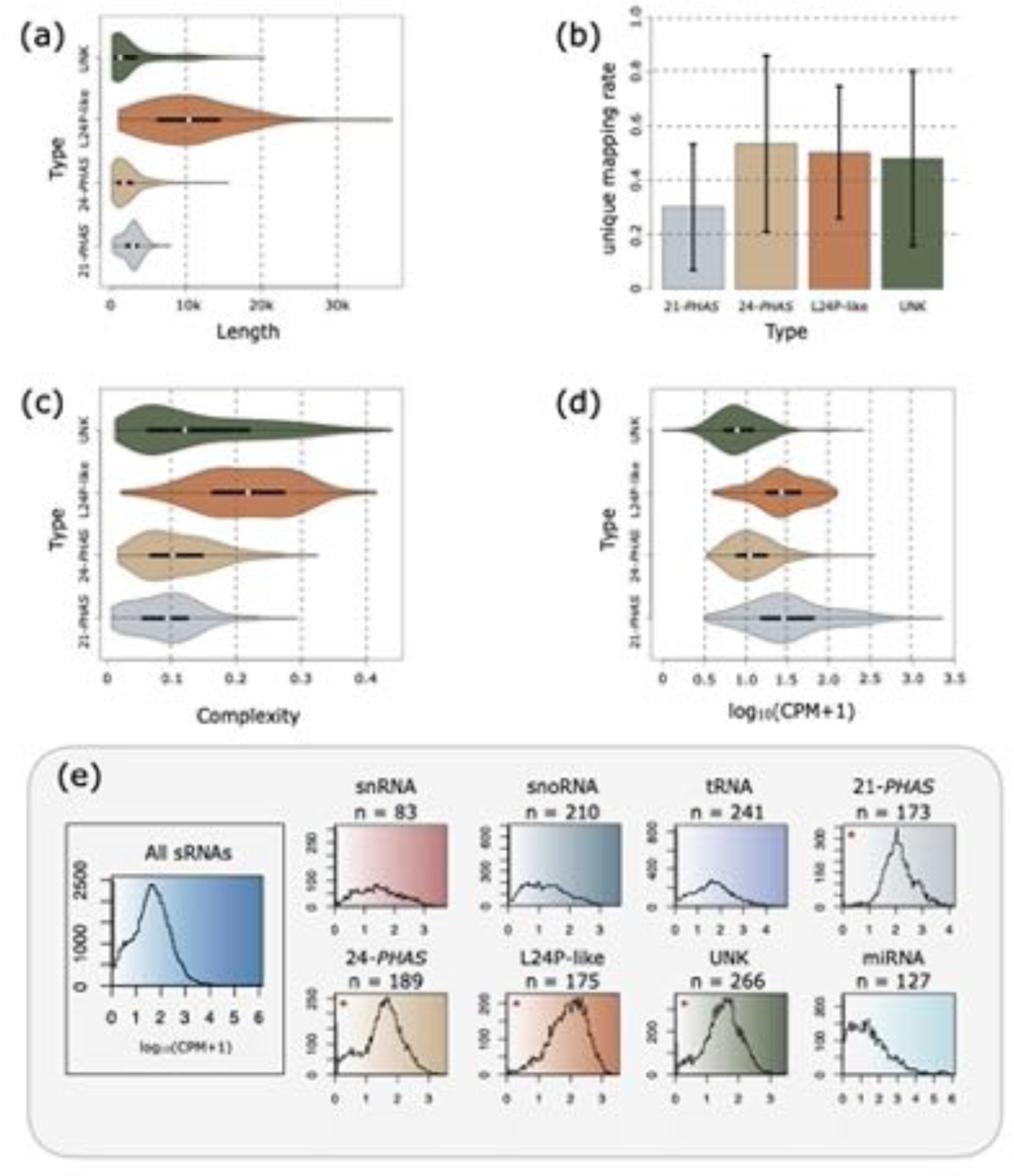
Overall features of phasing loci types in *Coffea arabica* during S0 (node) to flower development. (a) Length of different types of phasing loci displayed in kilobases. The L24P-like type contains the larger loci producing sRNAs. (b) The fraction of uniquely mapping reads aggregated across phased small RNA loci types. The *21-PHAS* sRNAs tend to, proportionally, map less uniquely than the other types. (c) Complexity of different types of phasing loci. The complexity parameter is calculated between 0 to 1 where 0 represents a less diverse set of sRNA fragments and 1 a more diverse set of fragments mapped to a loci. The *L*24P-like type has enhanced complexity compared to other types meaning that this type produces more diverse sRNAs. (d) Aggregated abundance across different types of phasing loci displayed as logarithmic base 10 of Counts Per Million (CPM) plus one. The *21-PHAS* is the type with most sRNAs fragments mapped to them while the UNK is the less abundant. (e) Abundance histogram (with 100 bins) of the different types of expressed sRNA producing loci displayed as logarithmic base 10 of Counts Per Million (CPM) plus one. Leftmost histogram shows the combined histogram of all expressed sRNAs loci (n = 1,462). Histograms marked with an asterix (*) shows loci with phasing patterns. (a,b,d) Boxes in the violin plots represent the interquartile range (Q1 to Q3) while the white circles represent the median and whiskers are set to 1.5 times the interquartile range. Maximum and minimum values are delimited at the extremities of the kernel density plots.

About 270 loci identified as phasiRNA-producing by ShortStack didn’t pass our manual evaluation (Table S5). Various reasons led us to exclude these loci as genuine phasiRNAs—such as low phasing scores (<15), presence in lengthy repetitive regions, strand biases, or low siRNA production, among others. These sequences were categorized as unknown (UNK), despite displaying phasing patterns.

Finally, for consistent L24P-like designation, we suggest that long 24-*PHAS*-like loci should possess a phasing score exceeding 15 (computed using Chen et al.’s predictive algorithm (Chen *et al*., 2007)) for 24-nt interval phasing and extend beyond 1,800-nt in length. We observed that L24P-like loci exhibited higher complexity compared to other phased loci (Fig. 2c), indicative of a wider array of sRNA fragments and less distinct phasing patterns. Despite being longer than other phased loci, L24P- like loci showed lower accumulation than 21-*PHAS* loci, yet higher accumulation than 24-*PHAS* and UNK loci.

### The flower transition is accompanied by an extensive reprogramming of sRNAs

The shift from vegetative to reproductive stages in coffee meristems, along with subsequent branch senescence, occurs biannually (Camargo & Camargo, 2001). Over these two years, various regulatory networks must decipher both internal and external signals to orchestrate the transformation of meristems at nodes from vegetative to floral stages, enabling multiple floral meristems induced at varying times to synchronize anthesis (Cardon *et al*., 2022). To delve into the roles of sRNAs, we sequenced and scrutinized 250 million sRNAseq reads obtained from eight developmental stages, spanning from nodes with undetermined vegetative growth buds (S0) to anthesis (detailed in Table S1). All examined reads were mapped to a curated reference.

Comparing sRNA abundance between S0 (nodes with buds under undetermined vegetative growth) and S1 stages revealed only 18 Differentially Accumulated (DA) sRNA loci. This indicates that S1 (node containing swollen buds) remains in a vegetative stage. However, contrasting sRNAs in S0 samples with more advanced stages (S2 to Flower; reproductive) revealed distinct patterns (Fig. 3, Dataset S1).

**Figure 3.**
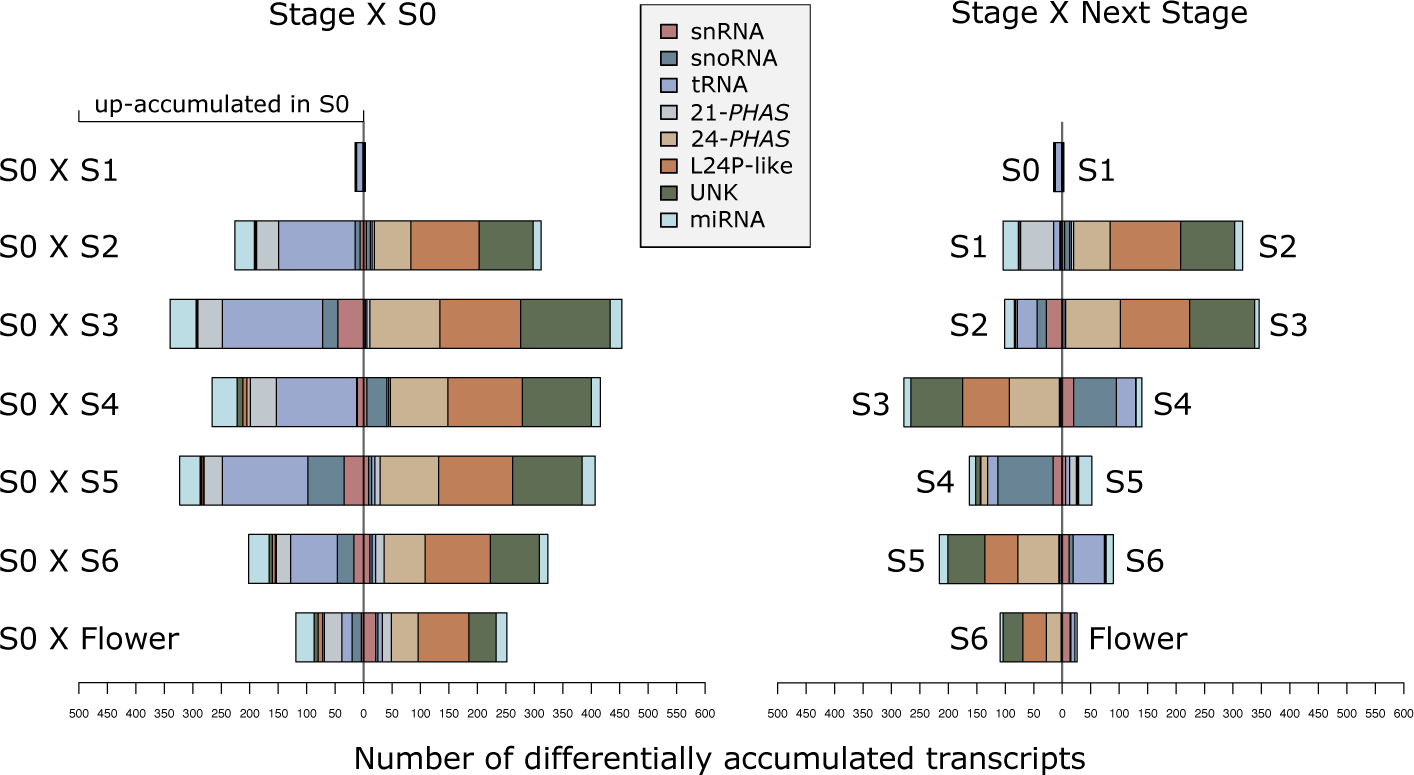
Number of differentially accumulated transcripts based on the small RNA sequencing quantification. Colors represent RNA types; bar represents the number of up accumulated transcripts in a given pairwise contrast between development stages. Left side summarizes the contrast of all stages against S0. Right side summarizes the contrast of a given stage against its subsequent stage. S0; node (with buds) undetermined vegetative growth. S1; Node containing swollen buds. S2; buds <3 mm. S3; Buds >3 mm and <6 mm - early stage. S4; Buds >3 mm and <6 mm - late stage. S5; buds from 6 to 10 mm, light green color. S6; buds >10 mm, white color. Flower; flowers after anthesis.

Firstly, we found that tRNAs are preferentially accumulated in the vegetative S0 stage, and the same pattern is followed by the 21*-PHAS* loci (Fig. 3, left side). On the other hand, 24*-PHAS*, L24P- like, and UNK are preferentially accumulated in the reproductive organs (S2, S3, S4, S5, S6 and Flower). As described above, some of the *PHAS*-like loci (L24P-like and UNK) may be siren RNAs (Rodrigues *et al*., 2013; Grover *et al*., 2020) or paternal epigenetically activated small interfering RNAs (easiRNAs) (Martinez *et al*., 2018) because of their accumulation in reproductive tissues.

Although consistently DA in reproductive samples, those 24*-PHAS* and 24*-PHAS-like* loci show a progressive reduction in accumulation after their peak in the S3 (Fig. 3, right side; Fig. S5). Contrarily, during the S3 to S4 transition, there was a noticeable rise in the accumulation of snRNAs, snoRNAs, and tRNAs (Fig. S6), peaking during S4 (Fig. 3, right side). Subsequently, a sharp reduction occurred in S5. Following this, snRNA and snoRNA levels remained steady, whereas tRNAs exhibited a preference for S0 and substantial reduction by S3. Some tRNA levels then surged after S4. Overall, our analysis highlights a distinct trend in tRNA, snRNA, and snoRNA accumulation, diverging from the patterns observed in 24-PHAS, L24P-like, and UNK. An interesting shift occurs during the S3 to S4 transition, commonly perceived as a single stage, denoted as G4 (Morais *et al*., 2008).

### The balance of miR156 and miR172 suggests juvenility restoration in S4

Two distinct groups of miRNAs exhibited preferential accumulation in vegetative or reproductive stages. The initial group was more abundant in the non-reproductive S0 and S1 stages, comprising 18 miRNA families such as miR164, miR169, miR171, miR172, miR319, miR394, miR396, miR399, and two putative novel miRNA families (Fig. S7). The second group consisted of four miRNA families: miR156, miR171, and two novel miRNA families. Their accumulation was notably higher in later stages like S5 or S6 (Fig. S8).

We observed a decrease in miR156 family abundance during stages S0, S1, and S2. However, certain miR156 family members showed preferential accumulation from S3 to flower stages, with a peak in S4. Conversely, miR172 family members exhibited an opposing pattern, being more abundant in S0, S1, and S2 (Fig. S9).

In Arabidopsis gametophytes, it has been demonstrated that miR156 members are reactivated *de novo* to reinstate the juvenile phase in each generation (Gao *et al*., 2022). The activation of miR156 members in S3, followed by a peak at S4, aligns with the understanding that these stages coincide with microsporogenesis and gametogenesis processes (Pimenta de Oliveira *et al*., 2023). The decreased accumulation of miR172 family members in S3 and S4 is in agreement with the finding that the floral induction takes place early in individual buds (Cardon *et al*., 2022) – probably before S2. Upon the establishment of an inflorescence meristem within a bud, as it reaches the S3 stage, it remains latent while other buds are being formed. Nonetheless, coordinated development persists until anthesis (Cardon *et al*., 2022).

### Co-expression analysis reveals that, besides miRNAs, sRNA loci of the same type are selectively co-regulated

To explore the accumulation patterns of distinct sRNA types from S0 to anthesis, we conducted a weighted gene co-expression network analysis using the WGCNA package. This analysis yielded five co-expression modules, labeled with Roman numerals I to V (Fig. 4a, Table S6). Each module primarily comprises a specific sRNA type: tRNAs, snoRNA, 21-*PHAS*, or 24-*PHAS*/L24P-like/UNK. In each module, we identified an eigen-gene representing a single transcribed element (edge), summarizing the module’s regulatory trend via Euclidean mean (Newman, 2006). In addition, a filtering step selected the most co-regulated members in each module by setting an adjacency threshold above the 95th quantile (Dataset S2).

**Figure 4.**
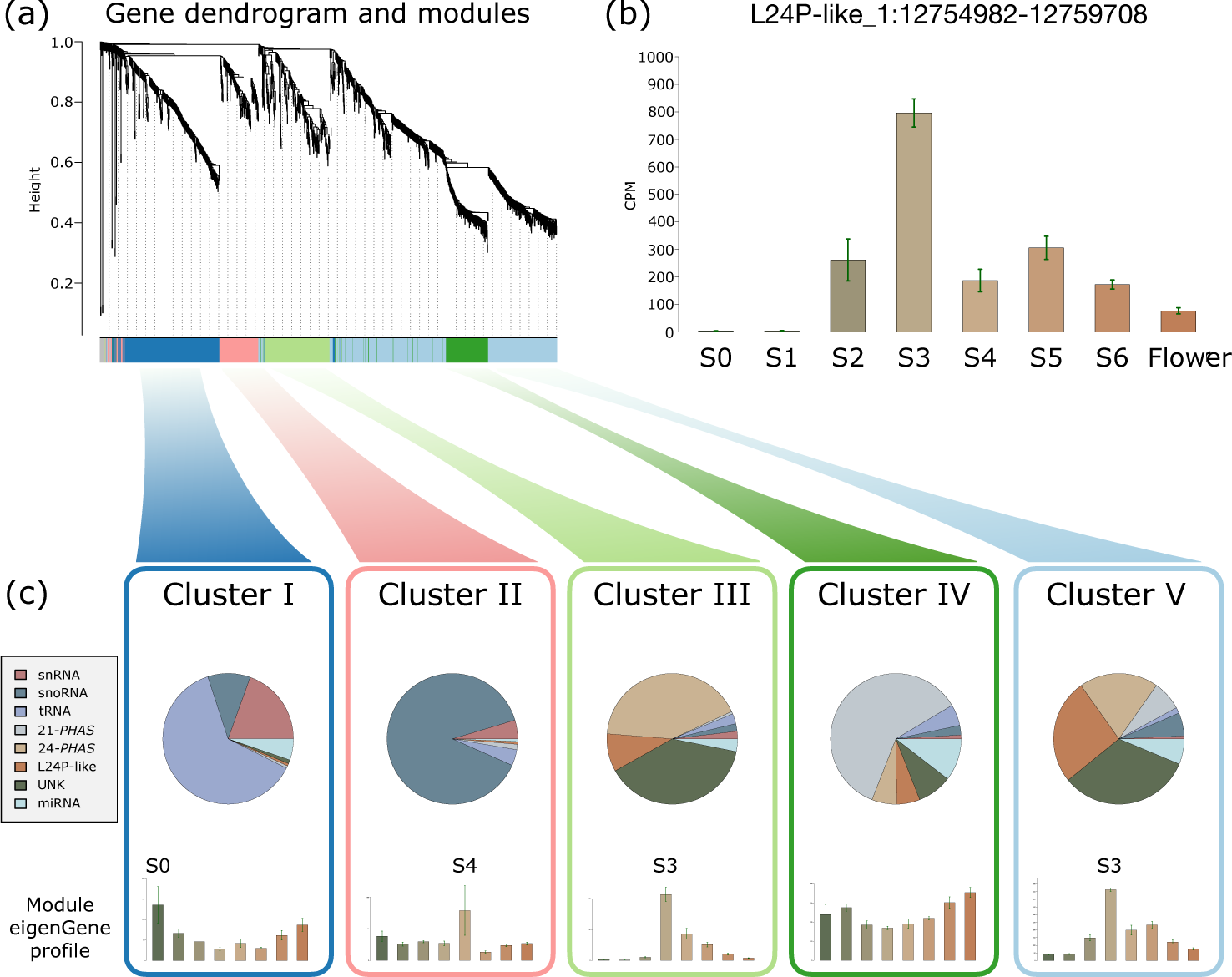
Co-expression analysis shows different types of sRNA producing loci are accumulated in a stage-specific way. (a) Gene dendrogram obtained by average linkage hierarchical clustering. The color row underneath the dendrogram shows the module assignment determined by the Dynamic Tree Cut. Different types of small RNAs producing loci are clustered based on their accumulation profiles. (b) Example of a *L24P- like* locus among the top connected nodes of Cluster V module showing its characteristic low accumulation level in S0 and S1, an increase in S2 and its peak in the S3 followed by a reduction onward. (c) Co-expression module composition (pie charts) and barplot representation of its eigen-gene in the stages, from left to right, S0, S1, S2, S3, S4, S5, S6 and Flower. Cluster I; composed mostly by tRNAs with expression peak in S0. Cluster II; composed mostly by snoRNAs with expression peak in S4. Cluster III; composed preferentially by *24-PHAS* and a significant proportion of *L24P-like* loci and UNK. This module is characterized by an expression peak in S3. Cluster IV; composed mostly by *21-PHAS* loci with the majority of members preferentially accumulated in S0 and S1, although many members are also constitutively present in all evaluated stages and some are accumulated in S6 and Flower. Cluster V; Similar but larger than Cluster III. This module is composed mostly by *24-PHAS*, *L24P-like* and UNK loci with expression peak in S3. However, contrary to Cluster III, this module displays an enhanced proportion of *L24P-like* loci.

The most extensive co-expressed module, Cluster V, comprises 578 members. Predominant within this module are 24-*PHAS/PHAS-like* loci, especially the unknown type (UNK). Its eigen-gene is a L24P-like with peak accumulation at the S3 stage. Employing an adjacency cutoff via the Topological Overlap Matrix (TOM), we identified highly co-regulated nodes representing miRNA families miR171, miR396, miR156, miR319, miR399, and three candidate novel miRNA families. Most elements within this module exhibit low accumulation in S0, S1, and S2, but experience a substantial increase in S3 (Fig. 4C). Cluster III exhibits a parallel trend to Cluster V, primarily composed of 24-*PHAS/PHAS-like* loci, albeit with fewer members (165). Its eigen-gene corresponds to the UNK type, peaking in S3. It made a negligible contribution in S0, S1, and S2 (Fig. 4c).

Cluster IV predominantly consists of 21-PHAS loci (∼60% of its 216 members). After filtering for top connected nodes, the proportion of 21-PHAS loci increased to 82%. Furthermore, miRNAs from families miR156 and miR482 were among the leading connected vertices within this regulatory module. We also detected four valine tRNAs among these highly connected nodes. Despite no distinct accumulation peak, module members remain relatively stable across stages, with a decrease apparent in S3 (Fig. 4c). Cluster IV exhibited the highest adjacency between its members, implying a finely-tuned accumulation profile. Co-regulation within Cluster IV remains relatively constant from S0 to Flower (Fig. 4c). This enduring regulatory control may stem from the significant proportion of disease resistance gene transcripts processed into 21-nt phasiRNAs, predominantly targeted by the highly abundant miR482 family. The sustained investment in transcriptional defense responses appears crucial to plant regulatory mechanisms. Consequently, tight control over the expression of defense genes could hold pivotal significance for *C. arabica* fitness.

Concerning Cluster I, the majority of its members are tRNAs (60% of 336 vertices), accompanied by a significant proportion of snRNAs and snoRNAs (19% and 10% respectively). However, after filtering for top connected vertices, the tRNA proportion rises to 92%. The module eigen-gene is a tryptophan tRNA with predominant accumulation in S0 and Flower stages (Fig. 4c). The accumulation profile of Cluster I members appears to align with that of Cluster IV, differing from the opposing trends observed in Cluster V and Cluster III.

Cluster II is primarily composed of snoRNAs (89% of 154 members), increasing to 93% for top connected vertices. Notably, module members, particularly snoRNAs, show preferential accumulation in S4 (Fig. 4c, Fig. S6). This distinct pattern stands in contrast to other modules. We hypothesize that this exclusive S4 peak may reflect active RNA metabolic processes and ribosome synthesis pathways, triggered as development resumes in response to water availability after a deficit.

### The transition from S4 to S5 coincides with shifts in the accumulation levels of miRNAs and their corresponding target genes

The transition from S4 to S5 is a pivotal phase in *C. arabica* flowering. This shift denotes the transformation from seemingly latent buds in S4 (buds >3 mm and <6 mm) to an active and rapidly progressing stage in S5 (buds ranging from 6 to 10 mm, displaying a light green color) (Majerowicz & Söndahl, 2005; López *et al*., 2021). We propose that S4 signifies the priming of the molecular machinery, facilitating the restart of reproductive development and the initiation of anthesis. To better investigate this transition, we produced RNAseq libraries from S4 and S5 stages as well as PARE libraries of samples from S4, S5 and anthers in pre-meiotic, meiotic and post-meiotic stages. The miRNA target prediction rendered a total of 3,213 miRNA-genic target pairs (Table S7). We also identified 2,003 inter-genic miR-target pairs in the *C. arabica* whole genome (Table S8). Fig. 5 summarizes the main findings regarding the accumulation levels of miRNAs and their target genes.

**Figure 5.**
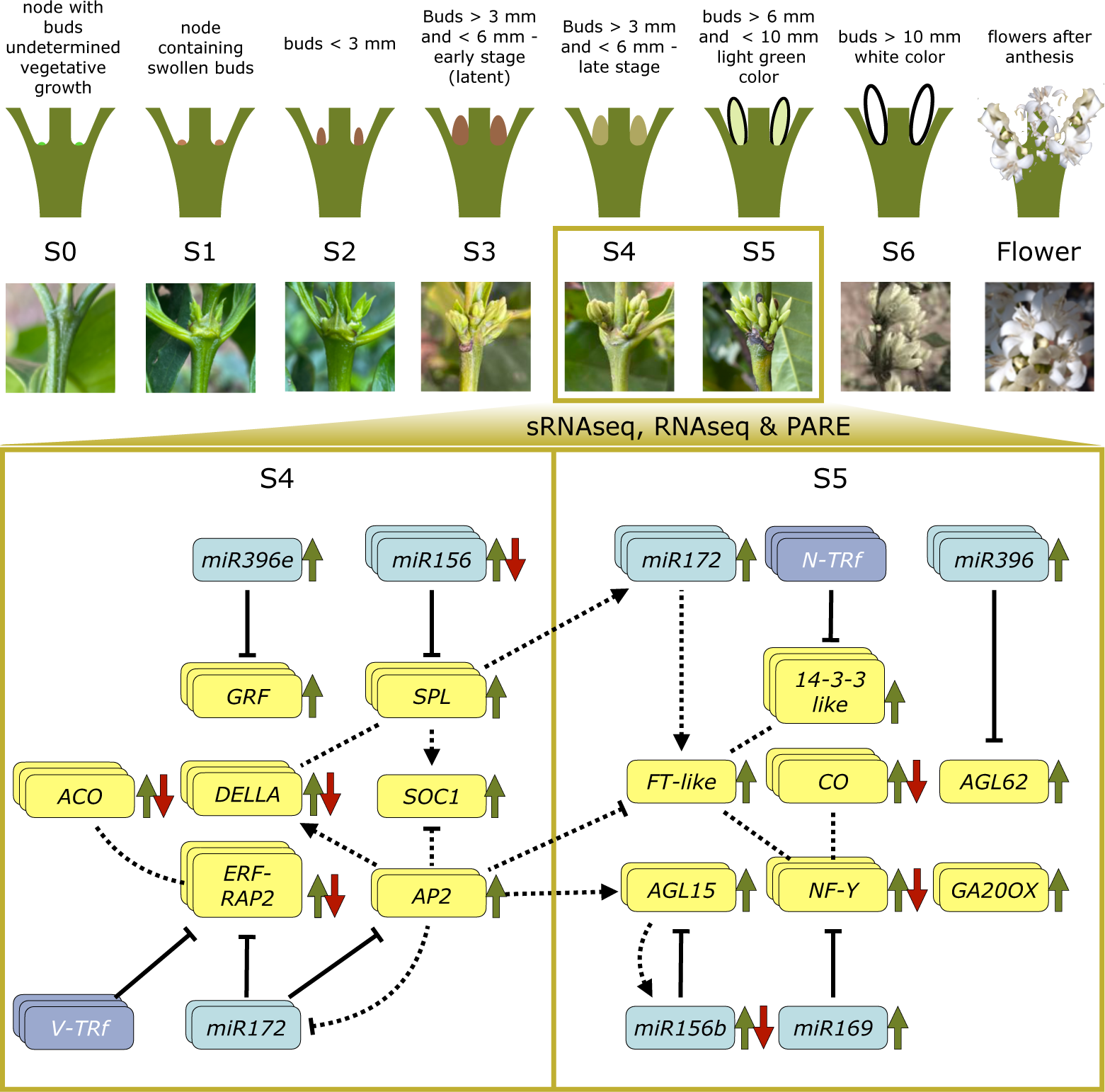
Schematic overview of the S0 to flower development and the core regulatory network of miRNA/t-RNAs and Protein Coding Genes governing the S4 to S5 transition. The combination of sRNAseq, RNAseq and PARE analysis reveals conserved motifs of gene regulation. The divided rectangle depicts the contrasting transcription profile between S4 and S5. Blue boxes represent *MIR* genes while Darker blue represents tRNA fragments (tRFs), yellow boxes represent Protein Coding Genes (PCGs). Single boxes show that a specific gene is being transcribed while two stacked boxes show that two homologs of that gene are transcribed and three stacked boxes show that three, or more, homologs of are transcribed. Green arrows pointed upwards represent genes being statistically more expressed in the S4 or S5 stage - according to the side which a gene is. Green and red arrows in opposite directions show that different homologs - or family members - of a gene present a contrasting transcription trend. Continuous black arrows with blunt ends between an interfering RNA and a PCG shows that the targeting was verified using PARE data. Dotted lines without arrows show that there is literature support of protein-protein interaction. Dotted lines with arrows between two genes shows that there is literature support that one gene is a direct activator of the other gene in other plant species.

### The miRNA families miR396, miR156 and miR172 are master modulators of PCG in S4 and S5

The most up-accumulated microRNA in S4 compared to S5 is a member of family miR396 (ccp- miR396e-3p) which targets ten *GROWTH-REGULATING FACTOR* (*GRF*) loci (Table S9). In grapevine, mutations in the miR396 binding site of *GRF4* caused changes in the inflorescence bunch architecture (Rossmann *et al*., 2020). In our PCG differential expression analysis, all miR396-targeted GRFs exhibited notably higher abundance in S4 compared to S5 (Table S10). While the most abundant MIR gene in S4 belongs to the miR396 family, it’s worth mentioning that certain family members also display increased levels in S5. Whereas the increased-in-S4 ccp-miR396e-3p targets *GFRs,* the group of loci encoding for three mature miRNAs increased in S5 are targeting genes such as *TIC110- chloroplastic*, *THREONINE-TRNA LIGASE,* and *AGAMOUS-LIKE MADS-BOX AGL62*. Of these target genes, only *AGL62* was found to be Differentially Expressed (DE), being up-regulated in S5. In *A. thaliana*, *AGL62* is only expressed in seed endosperm, regulating cellularization and acting as an upstream activator of *InvINH1*, an endosperm-specific invertase inhibitor (Kang *et al*., 2008; Hoffmann *et al*., 2022). In *C. arabica*, we identified the gene SubC.e_6694 – characterized as a putative invertase inhibitor – as up-regulated in S5. This miR396-AGL62-InvINH1 regulatory network may play a vital role in managing coffee floral bud growth rate by modulating energy (sucrose) levels as required.

Five members of family miR156 were more accumulated in S4 than S5 and we found them to be targeting seventeen *SQUAMOSA PROMOTER-BINDING-like* (*SPL*), preferentially at the pre meiotic stage. Only one *SPL* was up-regulated in the stages S4 compared to S5 (Tables S9 and S10). In Arabidopsis leaves, the interaction between the miR156-*SPL3* module and *FLOWERING LOCUS T (FT)* is part of the regulatory mechanism controlling flowering time in response to ambient temperature (Kim *et al*., 2012). Here, a FT-like gene exhibited up-regulation in S5 compared to S4. Conversely, the majority of miR156 family members displayed decreased accumulation in S5, and a SPL gene was also downregulated in S5.

Among the mature miRNAs more abundant in S5 than S4, we found a single member of miR156 family. Like the group of mature miR156 that accumulate in S4, this S5-enriched copy also targets *SPL* transcripts (Fig. S10, Table S11). This targeting of *SPLs* might function to modulate the activity of *FT* during S5 to synchronize floral development. The up-regulation of *AGL15* in S5 may explain why this miR156 member is more accumulated in libraries of later developmental stages such as S5, S6 and Flower.

### Ethylene responsive genes, such *AP2*, are DE during the transition of S4 to S5 and are target by miR172 family members

PARE analyses revealed two mature miR172 family members, miR172.2.ab, targeting four *ETHYLENE-RESPONSIVE TRANSCRIPTION FACTOR RELATED TO APETALA2-7-like* (*ERF-RAP2-like*) genes, predominantly in S4, with some targeting also observed in S5 (Fig. S11, Table S7). Notably, miR172.2.ab wasn’t found to be differentially accumulated between S4 and S5. However, one of its targets, APETALA2/ETHYLENE RESPONSIVE FACTOR (AP2/ERF), along with 41 other ERF genes, exhibited higher expression in S4 compared to S5 (Table S10).

This mature miR172.2.ab also targets two genes identified as “*floral homeotic protein APETALA 2”* (*AP2*) of which one was found to be up-regulated in S4. The Arabidopsis *AP2* promotes early flowering identity (Jofuku *et al*., 1994). It also has a subsequent function on the transition of the inflorescence meristem to a floral meristem (Drews *et al*., 1991) and plays a central role in the specification of sepals and petals (Krogan *et al*., 2012). *AP2* can also induce the expression of *AGL15*, a floral repressor, and directly down-regulate the transcription of floral activators like *SOC1* (Yant *et al*., 2010) and *FT* (Zhu & Helliwell, 2011). Here we found a contrasting expression profile of the *MADS-box SOC1* that is up-regulated in S4 while *FT* is up-regulated in S5 (Fig. 5).

### The enzyme responsible for the final step in ethylene biosynthesis is present in both S4 and S5 but the precursor step is missing

Among a myriad of biological processes, ethylene is also an important anthesis regulator in *C. arabica* (López *et al*., 2021). Ethylene biosynthesis involves two steps which a S-adenosyl-L-methionine (*SAM*) precursor is converted to 1-aminocyclopropane-1-carboxylic acid (*ACC*) by *ACC-synthase* (*ACS*) and then it is transformed into ethylene *by* 1-AMINOCYCLOPROPANE-1- CARBOXYLATE OXIDASE (*ACO*) (Zhang *et al*., 1995; Houben & Van de Poel, 2019). Nevertheless, the regulation of its biosynthesis is far more complex and occurs at multiple regulatory layers in specific tissues (Pattyn *et al*., 2021).

Among the 42 ERF genes up-regulated in S4 compared to S5, we identified 12 *ACO* showing increased expression in S4. Notably, the second and third most DE genes in S4 were *ACOs*. In contrast, ten other *ERF* and eight *ACO* displayed up-regulation in S5, but with a notably lower fold change (Table S10). Furthermore, no *ACS* exhibited differential expression between S4 and S5. These findings align with the notion that the conversion of SAM to ACC occurs in other tissues and is then transported to the inflorescence meristem, rather than being synthesized at the meristem itself (Lima *et al*., 2021).

### tRNAs are among the most up-accumulated sRNA genes in S5 and are putatively targeting PCGs and TE

The most highly DA sRNAs in S5 are four tRNAs, with their abundance in S5 being at least one hundred times greater than in S4 (Fig. S12). Additionally, the most abundant sRNAs in S5 were two lysine transporters, followed by an asparagine and threonine transporter. To ascertain whether these sRNAs play a regulatory role in flower development or are merely a by-product of heightened transcriptomic activity, we investigated further. Our analysis identified 1,373 tRNFs potentially targeting 1,131 PCGs (Table S12). Interestingly, we found six ethylene-responsive transcription factors predicted to be targeted by tRFs. In addition, two *14-3-3-like* transcripts - known to be intracellular receptors for the FT florigen in rice (Taoka *et al*., 2011) - were found to be targeted by a tRF from asparagine tRNA loci in meiotic and post-meiotic PARE libraries. Three *14-3-3-like* genes were found to be up-regulated in S5 shoot apical cells.

### The transcriptome of S4 exhibits hormone responsiveness and involvement in RNA biosynthesis, while that of S5 is primarily dedicated to cell wall production

snoRNAs and their associated proteins are ancient components that mediate maturation of ribosomal RNA (rRNA) (Bertrand & Fournier, 2013). The S4 and S5 stages mark an evident change in the accumulation of snoRNAs that are abundant in S4. In S5, there is a relative reduction in the levels of not only snoRNAs but also other sRNA types (Table S13, Fig. 3). In addition, a total of 39 ribosomal-related PCG were found to be DE between S4 and S5. The up-regulated ribosomal-related genes in S4 are predominantly coding for 60S ribosomal proteins and ribosome biogenesis proteins, whereas the up-regulated in S5 are 50S and 30S ribosomal protein components.

Because ribosomes are central to the protein translation pathways, we performed an overall analysis of GO terms of the 7,283 PCGs that were DE between S4 and S5 (Table S10). We classified those DE genes into 684 GO terms that revealed the development of male organs - with their respective meiosis processes, integration of hormonal signals, and intense metabolism of RNAs, in particular snoRNAs, are characteristic events of S4. Next, during S5, the expressed genes are mainly involved in processes to develop cell wall structures. The metabolism of structural carbohydrates, such as xyloglucan and xylan, is evident once in a matter of days the buds can grow from 6 mm to 10 mm, an increase of more than 60% in their length.

## Discussion

We conducted a genome-wide annotation of miRNAs and phasiRNA in the allotetraploid *C. arabica.* This data was then integrated with publicly available annotations of sRNAs (Johns Hopkins University, 2018), allowing us to investigate their abundance profiles along reproductive organ development. Finally, we coupled these datasets with RNAseq differential expression data from two key stages (S4 and S5) and PARE analysis.

We show that tRNAs are preferentially accumulated in S0 whereas 24-*PHAS* and *24-PHAS-like* loci are preferentially accumulated in S3 (Fig. 3 - left side, Fig. 4 - Clusters I, III, IV and V). The 24- *PHAS* accumulation coincides with an extended period of anatomical latency in which buds are apparently dormant - S3. This stage-specific accumulation also resembles the spatio-temporal dependent expression of many 24-nt phasiRNAs in species such as barley (Bélanger *et al*., 2020), wheat (Bélanger *et al*., 2020), maize (Zhai *et al*., 2015), tomato (Xia *et al*., 2019), petunia (Xia *et al*., 2019) and soybean (Arikit *et al*., 2014). Some of these 24-nt phasing sRNA loci could guide epigenetic imprinting, potentially including siren or easiRNAs (Polydore *et al*., 2018; Rodrigues *et al*., 2013; Grover *et al*., 2020; Martinez *et al*., 2018). Meanwhile, snRNAs and snoRNAs are preferentially accumulated in S4 (Fig. S6), in accordance with the most enriched GO term “RNA biosynthetic process”. We hypothesize that the re-activation of the transcription is an important feature for resuming flower development.

### The miR482 family is abundant in *C. arabica*, potentially for responses to pathogens

miRNAs, in accordance with their developmental regulatory role (Debernardi *et al*., 2022), were found to have families preferentially accumulated in vegetative or reproductive organs (Fig. 3 - left side; Fig. S7 and S8). However, miR482 superfamily members associated with resistance were consistently accumulated across all evaluated developmental stages in this study. miR482 superfamily members are widespread in most examined seed plants (Liu *et al*., 2020) emerging in Gymnosperms and latter functionally diversifying in eudicots and monocots (Xia *et al*., 2015). Here we found that miR482 is the family with more precursor loci in *C. arabica* - at least 20 - which is a similar number to Norway spruce with at least 24 loci (Xia *et al*., 2015). In addition, miR482 is the third most accumulated family.

The high number of miR482 loci, and the emergence of a putative novel mature miR482 in *Coffea* (*Novel-245889.2.ab*), supports the notion that there is a constant evolution of miRNA triggers of phasiRNAs that targets *R* genes (Liu *et al*., 2020). We hypothesize that this reflects an arms race between evolving components of plant resistance systems and evolutionary strategies to balance counter responses of potential and known pathogens (de Vries *et al*., 2015). *R* genes are among the first line of defense against biotrophic pathogen infection (Meyers *et al*., 2005). Their wide-spread in plant genomes allows specific resistance often associated with pathogen recognition followed by hypersensitive response in the form of localized programmed cell death (Heath, 2000; Meyers *et al*., 2003). Plant pathogens tend to infect specific hosts by complex interactions at the molecular level, which is a reflection of the host-pathogen coevolution (Meyers *et al*., 2005). We found that miR482 family members are targeting more than a hundred of putative *R* genes encoding for proteins containing nucleotide-binding site (NBS) and leucine-rich repeat (LRR) domains that were found to be 21-*PHAS*. We show that the Cluster IV was verified to have the highest mean co-expression of all modules (Dataset S2). These co-expression of miR482 and 21-*PHAS* are reflecting fine-tuned defense systems to provide proper control of responses to pathogens, probably avoiding mis-regulation of *R* genes.

### The S3, S4 and S5 transitions marks changes in sRNA abundance

The transition from S3 to S4 brings about significant changes in the accumulation of 24- PHAS/24P-like/UNK, snoRNAs, snRNAs, and tRNAs (Fig. 3), triggered by water availability following prolonged drought. Notably, in S4, the levels of 24-nt phasiRNAs decline noticeably, while there’s a substantial increase in structural RNA levels, particularly snoRNAs. This enrichment of snoRNAs defines Cluster II (Fig. 3 - right side, Fig. 4, and Dataset S2). This snoRNA accumulation is correlated with 60S ribosomal protein components that could regulate gene expression through enhanced translation efficiency (Kufel & Grzechnik, 2019).

The next transition, S4 to S5, is marked by a slow decrease in the accumulation of *24-PHAS* that will continue until anthesis (Fig. 3). Nevertheless, miRNAs levels increase in S5 suggesting tight regulation of developmental processes. Our PARE analysis showed that many of these miRNAs are targeting important genes related to flower development and response to ethylene (Fig. 5). These results support the role of ethylene as a key hormone in *C. arabica* flower development (Lima *et al*., 2021; López *et al*., 2021). We suggest that, as far as the transcriptome is concerned, the floral development is an event controlled at multiple levels that evolved to promote a synchronized anthesis in the face of endogenous and exogenous variability.

## Acknowledgements

We thank the Federal University of Lavras (UFLA/Brazil) and the members of the Laboratório de Fisiologia Molecular de Plantas (LFMP, UFLA/Brazil) for structural support of the experiments and analysis; Instituto Nacional de Ciência e Tecnologia do Café (INCT/Café), Coordenação de Aperfeiçoamento de Pessoal de Nível Superior (CAPES) and Conselho Nacional de Desenvolvimento Científico e Tecnológico (CNPq) for the financial support.

## Author contributions

A.C.J, B.C.M, C.N.F.B and R.R.O designed the study and conceptualized the project; T.H.C.R drafted the article, carried out bioinformatic analysis and design custom scripts; T.H.C.R and P.B carried out the phasing analysis; C.N.F.B and S.M performed data generation; T.H.C.R, T.C.S.C, M.S.G and L.R.A predicted *MIR* loci; C.N.F.B, K.K.P.O and G.L.R extracted RNA samples; A.C.J, B.C.M, P.B and R.R.O revised the manuscript.

## Data availability

Main data supporting the findings of this study are available within the paper and within its supplementary materials published online. Raw data for the RNAseq runs can be obtained through BioProject ID XXXXXXX. Raw data for the smallRNAseq runs can be obtained through BioProject ID XXXXXXXX - The respective SRA accessions are available in Supplemental Table S1. Raw PARE data can be obtained through BioProject ID XXXXXXX.

## Supporting information

Additional Supporting Information may be found online in the Supporting Information section at the end of the article.

**Dataset S1** Fasta sequences and differentially accumulated tables of sRNAs in coffea arabica vegetative to reproductive bud development.

**Dataset S2** Detailed information regarding the co-expression modules identified using the WGCNA package.

**Fig. S1** Cumulative abundance (in log of counts per million) of unique mature miRNAs in *C. arabica*.

**Fig. S2** Global multiple sequence alignment of mature miR482 family members.

**Fig. S3** *TAS3* and *TAS4* overall structure with their respective target sites.

**Fig. S4** Genome view of a Long 24-*PHAS*-like locus.

**Fig. S5** Abundances (counts per million) of representative 24-*PHAS*, L24P-like and UNK loci depicting their peak in S3.

**Fig. S6** Abundances (counts per million) of representative snRNA, snoRNA and tRNA loci.

**Fig. S7** Abundances (counts per million) of representative mature miRNAs from families preferentially up-accumulated in non-reproductive organs S0 and S1.

**Fig. S8** Abundances (counts per million) of representative mature miRNAs from families preferentially up-accumulated in late stages of flower development S4 to S6.

**Fig. S9** Abundances (counts per million) of mature sequences from families miR156 and miR172 in all libraries.

**Fig. S10** Mature miR156 preferentially accumulated in late stages of flower development S5 and S6.

**Fig. S11** Mature miR172 preferentially accumulated in vegetative stages S0 and S1.

**Fig. S12** Differential accumulation (depicted in counts per million) of four tRNA that are up- accumulated in S5 compared to S4.

**Table S1** Description of coffee bud stages during flower development based on transcriptional or phenological data.

**Table S2** 21-*PHAS* loci in the *C. arabica* genome.

**Table S3** 24-*PHAS* loci in the *C. arabica* genome.

**Table S4** Long 24-*PHAS*-like loci (L24P-like) loci in the *C. arabica* genome.

**Table S5** Unknown (UNK) loci with phasing pattern in the *C. arabica* genome.

**Table S6** Cluster members of co-regulated sRNAs.

**Table S7** miRNA genic targets in the C. arabica genome.

**Table S8** miRNA inter-genic targets in the *C. arabica* genome.

**Table S9** Up-accumulated miR in S4 and their target protein coding genes.

**Table S10** Differentially expressed PCG between S4 and S5.

**Table S11** Up-accumulated miR in S5 and their target protein coding genes.

**Table S12** tRNA derived fragments and their candidate target protein coding genes.

